# Hydrodynamics of linear acceleration in bluegill sunfish *Lepomis macrochirus*

**DOI:** 10.1101/386342

**Authors:** Tyler N. Wise, Margot A. B. Schwalbe, Eric D. Tytell

## Abstract

Bluegill sunfish accelerate primarily by increasing the total amount of force produced in each tail beat but not by substantially redirecting forces.

**ABSTRACT:** In their natural habitat, fish rarely swim steadily. Instead they frequently accelerate and decelerate. Relatively little is known about how fish produce extra force for acceleration in routine swimming behavior. In this study, we examined the flow around bluegill sunfish *Lepomis macrochirus* during steady swimming and during forward acceleration, starting at a range of initial swimming speeds. We found that bluegill produce vortices with higher circulation during acceleration, indicating a higher force per tail beat, but do not substantially redirect the force. We quantified the flow patterns using high speed video and particle image velocimetry and measured acceleration with small inertial measurement units attached to each fish. Even in steady tail beats, the fish accelerates slightly during each tail beat, and the magnitude of the acceleration varies. In steady tail beats, however, a high acceleration is followed by a lower acceleration or a deceleration, so that the swimming speed is maintained; in unsteady tail beats, the fish maintains the acceleration over several tailbeats, so that the swimming speed increases. We can thus compare the wake and kinematics during single steady and unsteady tailbeats that have the same peak acceleration. During unsteady tailbeats when the fish accelerates forward for several tailbeats, the wake vortex forces are much higher than those at the same acceleration during single tailbeats in steady swimming. The fish also undulates its body at higher amplitude and frequency during unsteady tailbeats. These kinematic changes likely increase the fluid dynamic added mass of the body, increasing the forces required to sustain acceleration over several tailbeats. The high amplitude and high frequency movements are also likely required to generate the higher forces needed for acceleration. Thus, it appears that bluegill sunfish face a tradeoff during acceleration: the body movements required for acceleration also make it harder to accelerate.

## INTRODUCTION

Many previous studies of the mechanics of fish swimming have focused on steady locomotion. But in nature, fish do not usually swim steadily. Instead, they rely on unsteady swimming maneuvers and changes in direction (Webb, 1991). Previous studies have examined some unsteady maneuvers like, C- and S- starts (Domenici and Blake, 1997), but few have looked at the hydrodynamics of linear accelerations. To better understand natural motion of fish, it is necessary to understand how fish accelerate. Similarly, in order to accurately replicate natural swimming motions in man-made objects, it is necessary to look at the forces during these natural behaviors.

The kinematics and hydrodynamics of acceleration in a variety of fish species were recently surveyed (Akanyeti et al., 2017). Across 51 species, they reported a consistent, large increase in tail beat amplitude and frequency. They also studied the wake of trout in detail, and showed that the kinematic changes lead to an increase in the impulse in vortex rings in the wake and a reorientation of the rings, indicating that the force is reorients to be more axially directed, rather than laterally. They found that the rings become more circular, which would result in a more hydrodynamically efficient transfer of force into the wake (Akanyeti et al., 2017). The hydrodynamics of acceleration in fishes has also been studied in American eels (Tytell, 2004a). Eels swim in an anguilliform mode, which means that they undulate most of their bodies during steady swimming and have a relatively short undulatory wavelength (Lauder and Tytell, 2005). During each tailbeat, they produce two pairs of vortices, a pattern called a 2P wake (Koochesfahani, 1989). The vortex pairs each produce a jet that is directed laterally, demonstrating that the eels have zero net force during steady swimming (Tytell and Lauder, 2004), as is required based on physical principles. During acceleration, the eels reoriented the vortex pairs so that the jets pointed away from the fish, more in the axial direction (Tytell, 2004a).

In this study, we examined the kinematics and hydrodynamics of acceleration in the bluegill sunfish, a carangiform swimmer that produces a different type of wake than the eel. Carangiform swimmers have longer body wavelengths than anguilliform swimmers and tend to have lower amplitude oscillations in the anterior body. During steady swimming, such carangiform swimmers produce a single pair of vortices per tail beat cycle (Drucker and Lauder, 2000; Lauder and Tytell, 2005; Tytell, 2007), a pattern called a 2S wake (Koochesfahani, 1989). In this type of wake, the jets between the vortex pairs point away from the fish even during steady swimming (Fig. 1A).

**Fig. 1:**
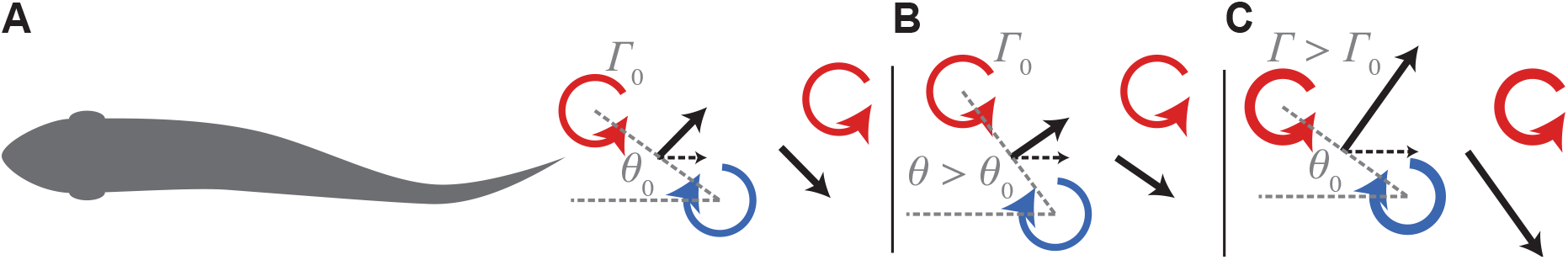
Schematic of hypotheses about how bluegill sunfish might produce more thrust during acceleration. Based on the steady swimming wake for a carangiform swimmer (A), the fish might produce more thrust by reorienting the tail force vector, which would be seen by an increase in *θ* for the vortex pairs in its wake (B), or by producing a larger force, which would be indicated by vortices with higher circulation *Γ* (C), or both. Red and blue circles indicate vortices, where *θ* is the angle of a vortex pair to the swimming direction and Γ is the vortex circulation. The thick black arrows and dashed arrows indicate total force and thrust force, respectively.

The steady wake structures of the bluegill and the eel suggest that they produce locomotor force in somewhat different ways. When a fish is swimming steadily, and acceleration is zero, the net force on the body should average out to zero over a tail beat cycle and over the entire body. This is equivalent to saying that the thrust force has to equal the drag force, on average. If the net force is zero, then there should not be any axial momentum in the fish’s wake. Using the wake to estimate the forces on a body in a fluid is a standard fluid dynamic technique, called a control volume analysis (Smits, 2000). For a steadily swimming eel, there is no net axial momentum in its wake, as would be expected by a control volume analysis (Tytell, 2007). In contrast, when a bluegill is swimming steadily, its wake contains a strong axial jet (Lauder and Tytell, 2005; Lauder et al., 2003; Tytell, 2007) (Fig. 1A). This might make it appear that thrust is not equal to drag during steady locomotion. This apparent discrepancy can be explained in several ways, as discussed in detail by Tytell (2007). First of all, a carangiform swimmer produces primarily thrust with its tail and primarily drag with its anterior body (Bale et al., 2014; Borazjani and Sotiropoulos, 2008; DuBois, 1978; DuBois et al., 1974). This spatial segregation of forces is similar to a boat with a propeller. An average over a full control volume would show a net zero axial momentum, but flow behind the hull will show net drag and flow behind the propeller would show net thrust. Similarly, for the bluegill, the wake close to the tail may mostly represent thrust from the tail (Tytell, 2007). Since eels do not segregate force production as much, the wake, even close to the tail, represents more of an overall average, and shows no net axial flow. Second, steady swimming is not actually steady; each tailbeat during steady locomotion is really a small acceleration and deceleration (Borazjani and Sotiropoulos, 2008; Tytell, 2007; Xiong and Lauder, 2014), and bluegill accelerate and decelerate more during steady swimming motions than eels (Tytell, 2007). The axial jets in their steady wake structure may represent the force needed for each small acceleration.

To accelerate, the fish must increase the axial force. Based on the wake structure during steady locomotion, there are two ways to increase the axial force generated. The fish could reorient the existing force, so that it points more in the axial direction (Fig. 1B), or it could increase the force, without changing the angle (Fig. 1C), or both. These two effects would cause characteristic changes in the wake. If the fish reorients the force, it will also reorient the vortex pairs in its wake, increasing the angle *θ* between the vortex pairs and the swimming direction, so that more of the resulting force is in the axial direction (Fig. 1B). For a bluegill, this angle is largely determined by the tail beat frequency, tail beat amplitude, and swimming speed. For example, if swimming speed and tail beat amplitude stay the same but tail beat frequency increases, then the vortices will be closer together, which would lead to an increase in the angle *θ*. If it increases the total force, it will produce vortices with higher circulation *Γ*. In this case, the force vector might point in the same direction as it does during steady locomotion, but the total force would be greater, meaning that the axial component of the total force is also larger (Figure 1C). Akanyeti et al. (2017) found that trout use both mechanisms; they produce more force and they reorient it more in the axial direction.

The increase in axial force must not only be large enough to accelerate the fish’s own mass, but it must also push the fluid ahead out of the way. This effect is called “added mass” (Faber, 1995); it is as if the fish had a larger mass than only its body. For a carangiform fish like the bluegill, the tail must also produce enough force to overcome the drag on the body (Tytell, 2007). The axial force that the tail produces during acceleration is thus composed of two parts: the force to accelerate the mass of the body and the fluid around it and the force to overcome the drag on the body.

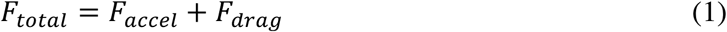

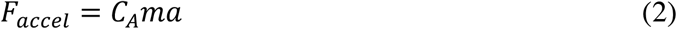

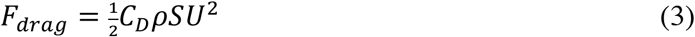

where *C_A_* is the dimensionless added mass coefficient, *m* is the mass of the body, *a* is acceleration, *C_D_* is a dimensionless drag coefficient, *ρ* is the density of water, *S* is an area (commonly the surface area of the body), and *U* is the swimming speed. The added mass coefficient *C_A_* is usually greater than 1, and both it and the drag coefficient *C_D_* may be different at different swimming speeds due to changes in tail beat frequency and amplitude or what fins the fish uses at different speeds.

Because bluegill are stiffer than eels or trout (Aleyev, 1977), we hypothesize that they may be less able than these fishes to achieve sufficiently high tail amplitudes to reorient vortices in the wake. Additionally, because bluegill have relatively long body wavelengths compared to these fishes (Lauder and Tytell, 2005), greater tail amplitudes may not lead to larger spacing between vortices in the wake, but may simply the head to swing from side to side more, an effect called recoil (Lighthill, 1970). Together, these two features of bluegill swimming suggest that they may accelerate differently than eels or trout, increasing the force from each tail beat (Figure 1C) rather than reorienting it.

In this study, we examined how bluegill sunfish produce forces during steady swimming and linear forward acceleration, which we term unsteady swimming. We quantified kinematics with high speed video, examined the wake using particle image velocimetry, and measured acceleration using an inertial measurement unit (IMU). For both steady and unsteady swimming, each tail beat contains forward acceleration (Tytell, 2007; Xiong and Lauder, 2014), which may be smaller or larger depending on the swimming mode and speed of the overall acceleration. This pattern allowed us to compare the forces and kinematics required for the same magnitude acceleration in isolated tailbeats during steady swimming and sustained over several tailbeats during unsteady swimming.

## MATERIALS AND METHODS

### Animals

Four bluegill sunfish (*Lepomis macrochirus* Rafinesque) were captured by beach seine in White Pond, Concord, MA, USA. All animals were housed individually in 38 l aquaria with 12 L: 12 D cycle and were fed live worms or flake food (earthworm flake, Pentair, Apopka, FL, USA) daily. Water temperature (20±2°C) and pH (7.4) were kept constant and were equal to that used during experiments. Fish total length ranged from 148 to 163 mm (mean ± s.d. = 155±7 mm) and mass ranged from 54 to 78 g (70±11 g). Animal care and all experimental procedures followed a protocol approved by Tufts University (M2012-145 and M2015-149).

### Particle image velocimetry

Flows generated by unsteady and steady swimming fish were quantified using particle image velocimetry (PIV) (Tytell, 2011). Neutrally-buoyant glass particles (50 *μ*m diameter, TSI Inc., Ypsilanti, MI, USA) were placed into a 293 l flow tank (Loligo Systems, Viborg, Denmark) and were illuminated with two 5W 532 nm continuous lasers (Opus 532, Laser Quantum, Santa Clara, CA, USA). One laser aimed directly at the viewing area in the flow tank (25×26×30 cm). The second laser projected a horizontal light sheet at a 45° mirror to increase the light intensity in viewing area (Fig. 2). The two light sheets were 6 cm from the bottom of the tank. Video was recorded from below the flow tank with a high speed camera (Phantom Miro M120, Vision Research, Wayne, NJ, USA) at 500 frames per second (Fig. 2).

**Fig. 2:**
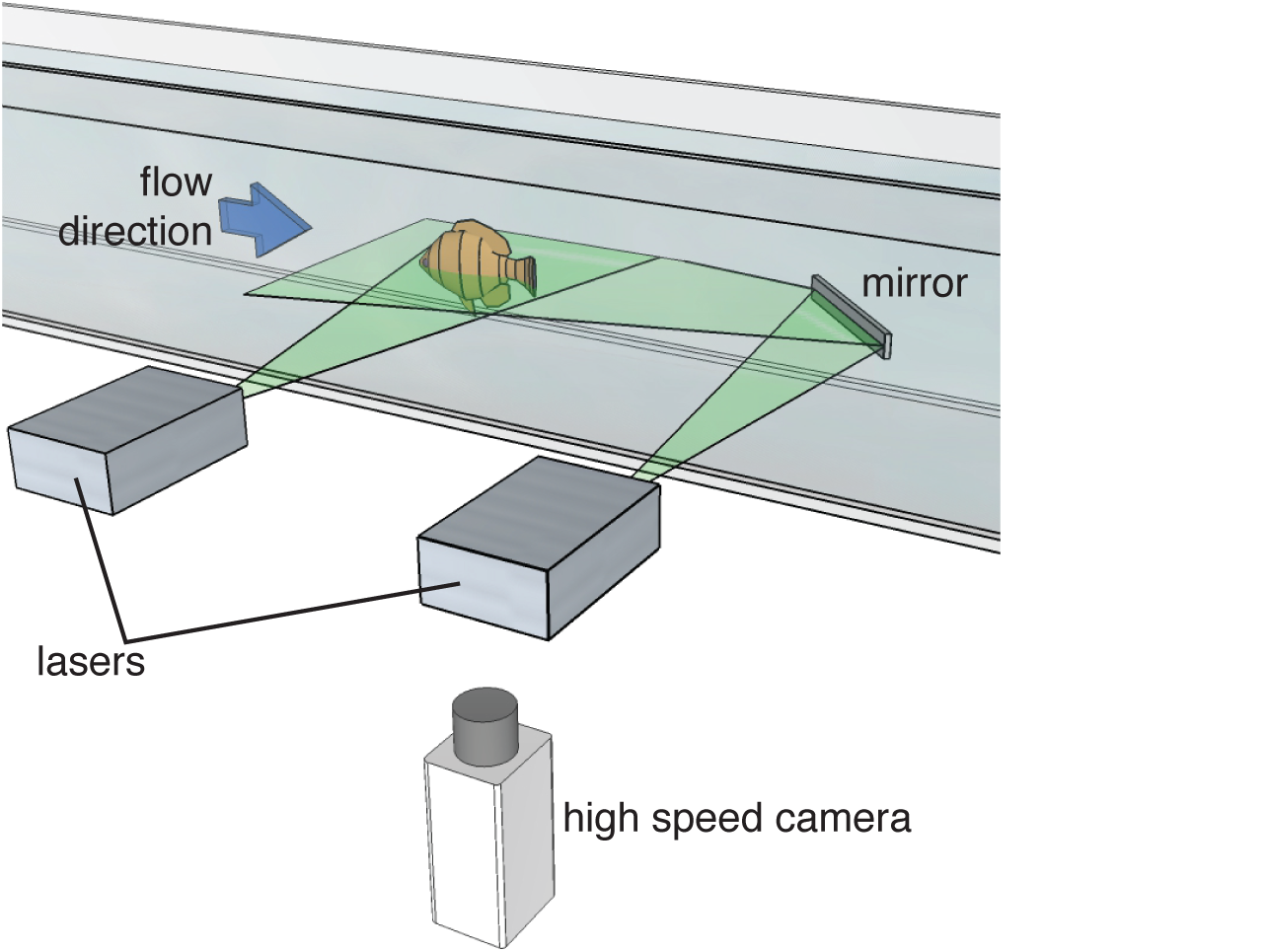
Experimental setup for visualizing flow during accelerations. A bluegill sunfish swims against the flow (blue arrow) in a flow tank. The fluid is illuminated by two lasers, one of which bounces off a mirror in the flow to illuminate the back side of the fish. A high speed camera below the tank records motion of particles in the flow.

### Inertial measurement unit construction and attachment

An inertial measurement unit (IMU; MPU-9250, InvenSense Inc., San Jose, CA, USA) was attached to the fish’s body to measure body orientation and dynamic acceleration. Fine, coated copper wires (80 μm diameter, Omega Engineering, Inc., Stamford, CT, USA) were soldered to individual pads on the IMU following the company’s instruction for the serial peripheral interface (SPI). The IMU and copper wire connections were waterproofed by encasing them in epoxy (CircuitWorks Epoxy, Chemtronics, Kennesaw, GA, USA). Data was collected from the IMU by connecting it to a USB SPI interface (USB-8451, National Instruments, Austin, TX, USA) and data was viewed in a custom LabVIEW program (v. 2014, National Instruments, Austin, TX, USA). Before surgery, in order to account for drift of the IMU data, the IMU was suspended in water with no flow for ten minutes. This was used to calibrate the drift during the trials.

An IMU was attached to each bluegill sunfish immediately before swimming experiments. Each fish was anesthetized with a buffered 0.02% solution of tricaine methane sulfonate (MS222, Sigma Aldrich, St. Louis, MO, USA). During surgery (~20 minutes), anesthesia was maintained by pumping buffered 0.01% MS222 over the fish’s gills. The IMU was sutured just below and anterior to the dorsal fin, which is near the fish’s center of mass, and a local anesthetic (bupivacaine USP, 1 mg/kg; Sigma-Aldrich, St. Louis, MO, USA) was injected in the suturing area.

Before the fish completely recovered from anesthesia, we calibrated the orientation of the IMU on the fish’s body. The fish was placed in three known orientations (left side down in a horizontal orientation, dorsal side up in a normal swimming orientation, and nose down in a vertical orientation) to identify the three axes in the coordinate system.

Fish recovered from the surgery in a separate tank before being placed in the flow tank (~10 min). Fish acclimated to the flow tank for an hour before experiments began. Upon completion of the experiments, fish were briefly anesthetized (0.01% buffered MS222) so that the IMU and sutures could be removed.

### Swimming Trials

Before an experiment could be conducted, each fish had to be acclimated to the flow tank and the laser sheets. Fish typically avoid the bright laser sheets and thus needed conditioning to acclimate to them. Fish were transported between their home tank and the flow tank to habituate to the experimental setup. Once the flow tank, fish swam into the flow and were gently prodded into the laser sheets by placing dowels in front and behind the fish. This procedure was repeated several times over four days until each fish would swim steadily within the light sheets for extended periods of time.

During experimental trials, each fish swam at flow speeds between 1.0 to 2.5 body lengths per second (BL s^−1^) in 0.5 BL s^−1^ intervals. Fish were confined to a 90 cm long section of the 25×26×150 cm (height × width × length) working section of the flow tank. For each bluegill sunfish, at least three acceleration trials and three steady locomotion sequences were recorded at each flow speed. Each trial had at least five tail beats, and the fish did not turn more than 30 degrees from its original position. During acceleration trials, we also recorded at least five tail beats, and all were analyzed.

We classified individual tail beats as “steady” or “unsteady” by measuring the motion of the fish in the laboratory frame of reference. When a fish is steadily matching the oncoming flow in the flow tank, its position is steady relative to the camera. A tail beat was classified as steady when a fish moved less than 2% of its body length over the course of the tail beat. Unsteady tail beats were those when it moved forward more than 2%, and we did not analyze tail beats in which the fish moved backward more than 2% of body length.

Accelerations were induced in multiple ways. Fish were first positioned in the viewing area using a pair of dowels. Accelerations were initiated by either removing the dowel in front of the fish or dropping a heavy object (e.g., a D-cell battery) behind the fish. Care was taken to avoid any hydrodynamic interference with either acceleration inducing method and the dowels. Fish occasionally responded with a C-start to the dropping of the heavy object, and these were not included in the analysis. During all experiments, fish were maintained in the center of the viewing area of the flow tank since swimming near the walls can increase thrust, based on measurements of flapping foils (Fernández-Prats et al., 2015).

### Data Analysis

Videos were processed with Insight 4G (TSI, Inc., Shoreview, MN, USA) to quantify fluid vorticity and velocity. The flow fields were processed using custom Matlab (R2014b, Mathworks, Natick, MA, USA) program. The center and diameter of vortices were manually identified and the circulation was calculated as

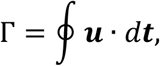

where ***u*** is the velocity vector and ***t*** is the unit vector tangent to the outline (Faber, 1995). The position data was used to calculate the orientation and distance in between each vortex pair. The total force, *F_mean_*, required to produce the vortex pair can be expressed by

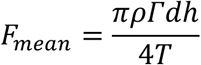

where *ρ* is the density of water, Γ is the average circulation of the two vortices, *d* is the distance between the two vortices, *h* is the height of the caudal fin, and *T* is the tail beat period (Tytell, 2011). The tail beat period for each individual tail beat was found using custom Matlab program to track the head and tail position of the fish over time. Force in the axial direction is thus

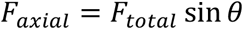

where *θ* is the angle made between two vortices and the forward motion of the fish (Fig. 1A).

The IMU measures angular velocities in three axes, and total acceleration along three axes. Total acceleration is the vector sum of dynamic acceleration vector due to the fish’s movement and the gravitational acceleration vector. To estimate just the dynamic acceleration, we use the algorithm developed by Madgwick *et al.* (2011) to estimate the orientation of the fish. Once the orientation is known, then the gravitational vector can be subtracted from the total acceleration to estimate dynamic acceleration. Briefly, the algorithm takes advantage of the fact that the IMU gives us two independent estimates of orientation. First, we can estimate orientation by integrating the angular velocities. This value tends to drift if angular velocities are low. But when angular velocities are low, dynamic accelerations also tend to be low, so that the total acceleration is close to the gravitational acceleration, which gives another estimate of orientation. By merging these two estimates in an optimal fashion, we can accurately estimate orientation (Madgwick et al., 2011) and from that estimate dynamic acceleration. The algorithm was implemented in a custom Matlab script.

A custom Matlab program allowed manual identification of the location of the tail tip and the tip of the snout. Based on these identified positions, the peak lateral excursion of the tail was identified, following procedures used previously (Tytell, 2004b). The tail beat amplitude is the distance from the midpoint to the peak excursion on either side. If *t_i_* is the time of peak lateral excursion and *t*_*i*–1_ is the time of the previous peak, then the tail beat frequency at time *i* is 1/2 (*t_i_* – *t*_*i*–1_). Head amplitudes were identified in the same way. Peak accelerations were extracted from the dynamic acceleration from the IMU, which determined the peak acceleration for each half tail beat, the time interval between peak lateral tail excursion on one side to peak excursion on the other (from *t*_*i*–1_ to *t_i_*).

### Statistics

Our statistical models were based on the physical model in Eqns. (1) – (3). We used a mixed model regression to compare how the total wake force depends on the swimming speed and acceleration. Although Eq. (1) represents a continuous relationship, neither total force nor acceleration were normally distributed, which would tend to result in biased or inaccurate regression coefficients. Total force was always positive, but had a long tail, making it appropriate for a log transformation. We then binned acceleration *a_dyn_* into four categories: zero (–1 ≤ *a_dyn_* < 1 BL s-2), low (1 ≤ *a_dyn_* < 2.5 BL s-2), medium (2.5 ≤ *a_dyn_* < 6 BL s-2), high (*a_dyn_* ≥ 6 BL s-2). The boundaries between bins were chosen so that we had approximately the same number of unsteady acceleration tailbeats in each bin. This procedure is similar to the ranking procedures that are the basis of most nonparametric statistics (Kloke and McKean, 2015). We also included the tailbeat type (steady vs. unsteady) and the flow speed in the model, along with the two-way interactions. Thus, the model for total force was

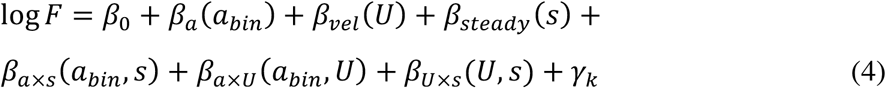

where *β*_0_ is the intercept, *β_a_*(*a_bin_*) is related to the added mass coefficient for each acceleration bin, and *β_vel_*(*U*) indicates the effect of flow speed on total force. The interaction terms describe how total force may depend on the combinations of acceleration, tailbeat type, and swimming speed. We included a random effect *γ_k_*, due to differences in individual fish.

We use the same model structure for vortex ring angle, diameter, tail beat frequency, and the head and tail amplitude. Tail beat frequency was also log transformed.

Regressions were performed in R (version 3.4.4, R Core Team, 2018) using the nlme package (version 3.1-131.1;Pinheiro et al., 2018). Marginal means were estimated using the emmeans package (version 1.2.1; Lenth, 2018). Figures were created using ggplot2 (version 2.2.1; Wickham, 2009) with colormaps from viridis (version 0.5.1; Garnier, 2018) and beeswarm plots with ggbeeswarm (version 0.6.0; Clarke and Sherrill-Mix, 2017).

Most figures use standard statistical box plots to show the distribution of the data, where the boxes span the 25th to the 75th percentile, with a line at the median, and whiskers represent 1.5 times the interquartile range.

## RESULTS

We tested the hypothesis that bluegill sunfish produce axial force by increasing the total amount of force produced, as opposed to reorienting the forces in its wake. Data was taken from four individuals at 1.0 BL·s^−1^, 1.5 BL·s^−1^, 2.0 BL·s^−1^, and three individuals at 2.5 BL·s^−1^. One individual would not swim steadily at 2.5 BL·s^−1^. A total of 1122 vortex pairs and the accompanying kinematics were analyzed.

Because we measured acceleration directly, we quantified acceleration in both steady tailbeats in which the fish matched speed against the flow and did not move more than 2% of its body length within the flow tank, and in unsteady tailbeats in which the fish moved forward in the flow tank. Even in nominally steady tailbeats, each tailbeat includes an acceleration and deceleration, but in unsteady sequences, the fish maintains the acceleration over several tailbeats, causing it to move forward in the tank. Fig. 3 shows the number of tailbeats in each acceleration category for steady and unsteady sequences.

**Fig. 3:**
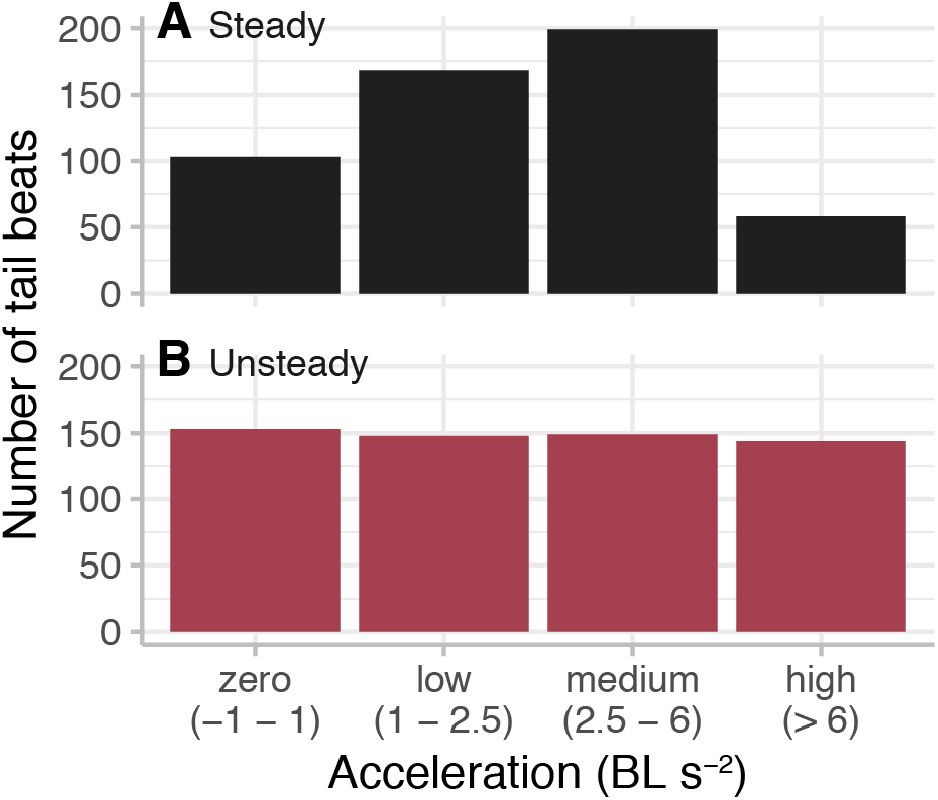
Number of tailbeats in each acceleration category. **A.** Steady swimming tailbeats. **B.** Unsteady tailbeats.

Figure 4 shows examples of the flow patterns in the wake during steady swimming with *a_dyn_* = 0.59 BL s^−2^ (Fig. 4A), and during two unsteady sequences, one with a medium accelerationwith *a_dyn_* = 4.75 BL s^−2^ (Fig. 4B), and the other with a high acceleration with *a_dyn_* = 67.7 BL s^−2^ (Fig. 4C),. The corresponding video sequences are available in Movie S1, S2, and S3. The wake consists of vortices that alternate in rotational direction, represented by red and blue colors on the figure. This created backward jets of fluid indicated by the velocity vectors, shown in black. Vorticity is higher and jets are stronger in the high acceleration sequence (Fig. 4C). The right panels of Fig. 4 show raw kinematic and acceleration data. Note that the acceleration (brown) has two peaks per tailbeat cycle, seen most clearly in Fig. 4A and B. This is expected because tail movements to the left side should produce a symmetrical forward acceleration as tail movements to the right. Higher accelerations correspond to increases in vortex circulation (red circles) and to changes in tail beat frequency. Additionally, the unsteady sequences (Fig. 4B and C; Movies S2 and S3) have higher tailbeat amplitude overall than the steady sequence (Fig. 4A; Movie S1), even most of the acceleration values in Fig. 4B are of similar magnitude to those in Fig. 4A.

**Fig. 4:**
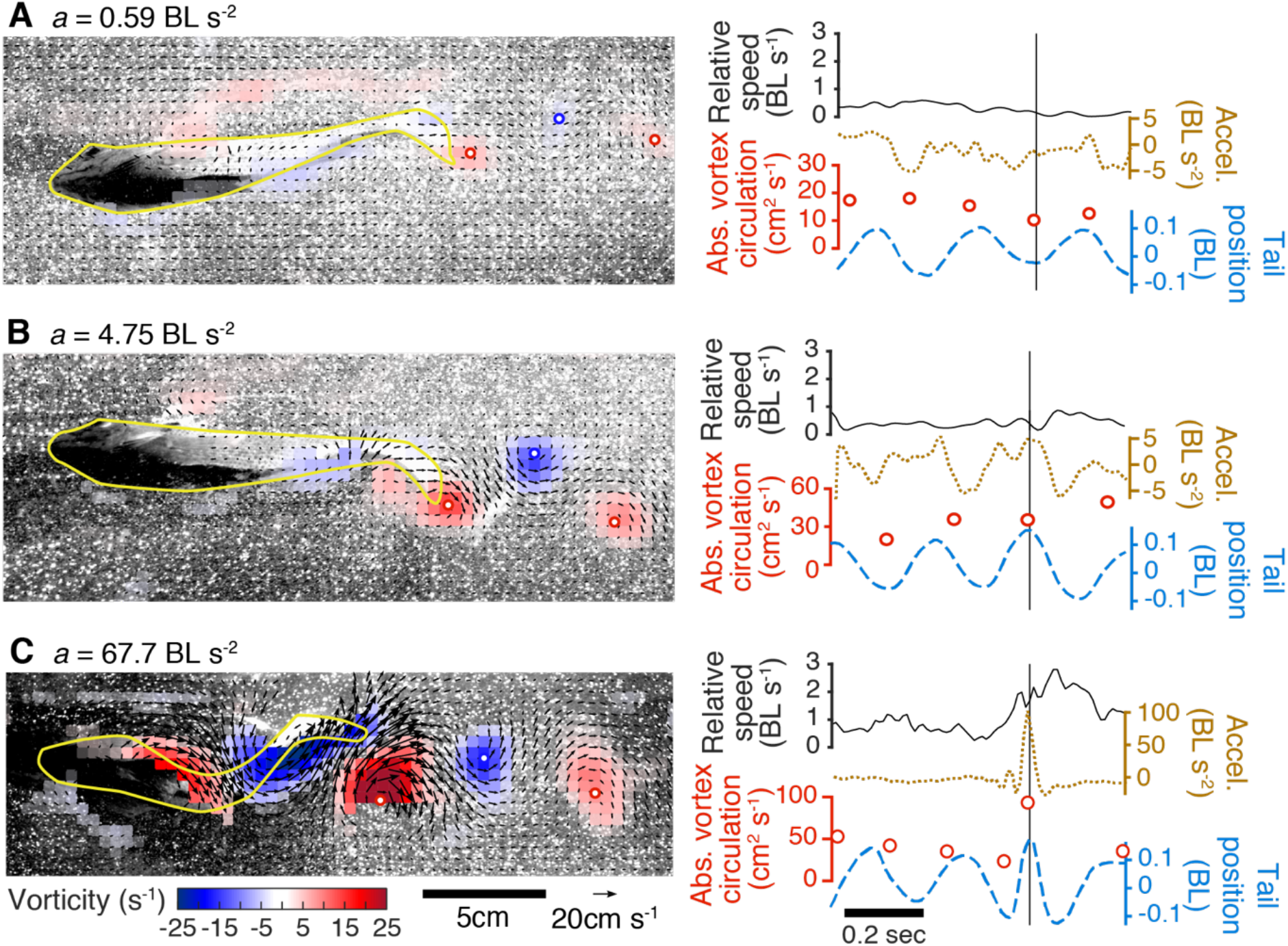
Example flow fields, kinematic data, and acceleration measurements. Ventral video frames, vorticity (in color), and velocity vectors (black arrows) for the same individual swimming steadily (A), and accelerating at a medium (B) and high rate (C), starting at an initial swimming speed of 1.5 BL s^−1^ in each case. Color indicates vorticity, with red and blue circles to indicate the centers of each vortex. The panels on the right show the speed relative to the laboratory (black), the dynamic acceleration (brown, dotted), the magnitude of the most recently shed vortex’s circulation (red circles), and the position of the tail (blue, dashed). The vertical line indicates the time of the frame shown on the left.

### Wake structure and force production

We compared the total force produced during steady and unsteady sequences with different magnitudes of acceleration, starting from different steady swimming speeds (Fig. 5). On average, force increases significantly in higher acceleration bins (p < 0.0001; Table 1), and is significantly higher in unsteady sequences compared to steady ones (p < 0.0001). Force also increases across swimming speeds (p < 0.0001), with the force at each flow speed significantly different from each other speed. The forces for unsteady high acceleration tailbeats are at least 2.24 times the forces for steady zero acceleration tailbeats (at 1.5 BL s-1) and are as much as 7.4 times (at 1.0 BL s-1) (Fig. 5).

**Table 1.**
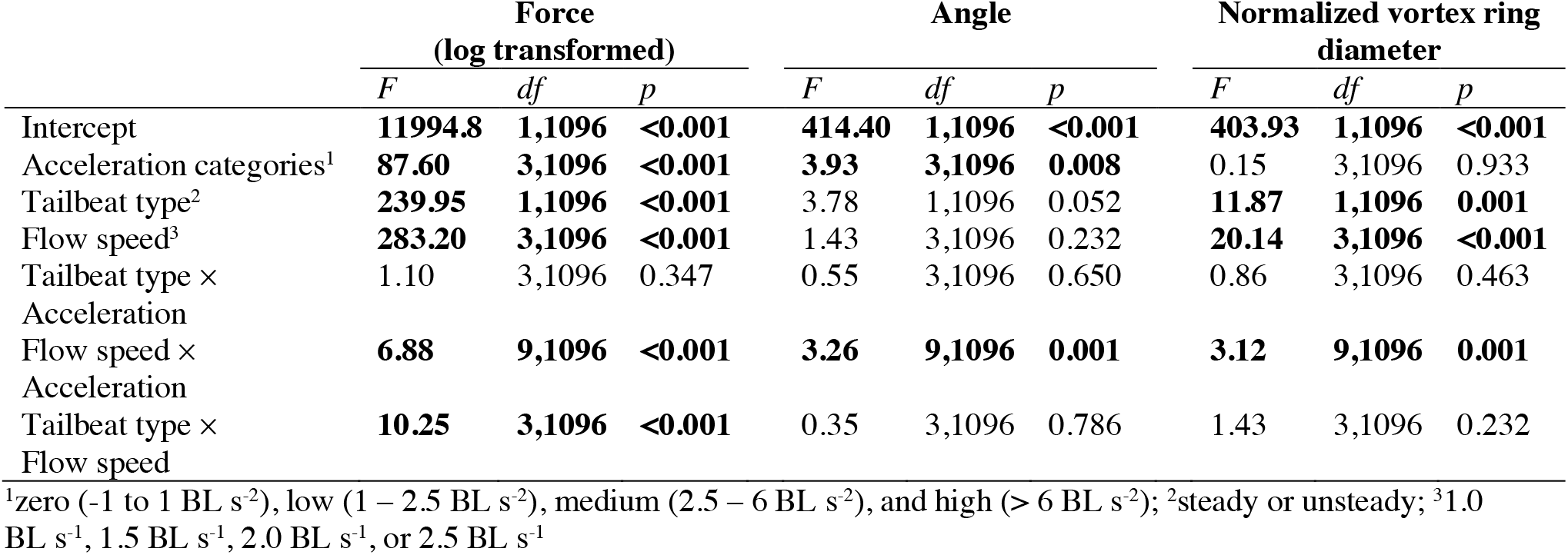
Mixed model regression results for relative force (log transformed) and wake vortex angle, and normalized vortex ring diameter.

**Fig. 5:**
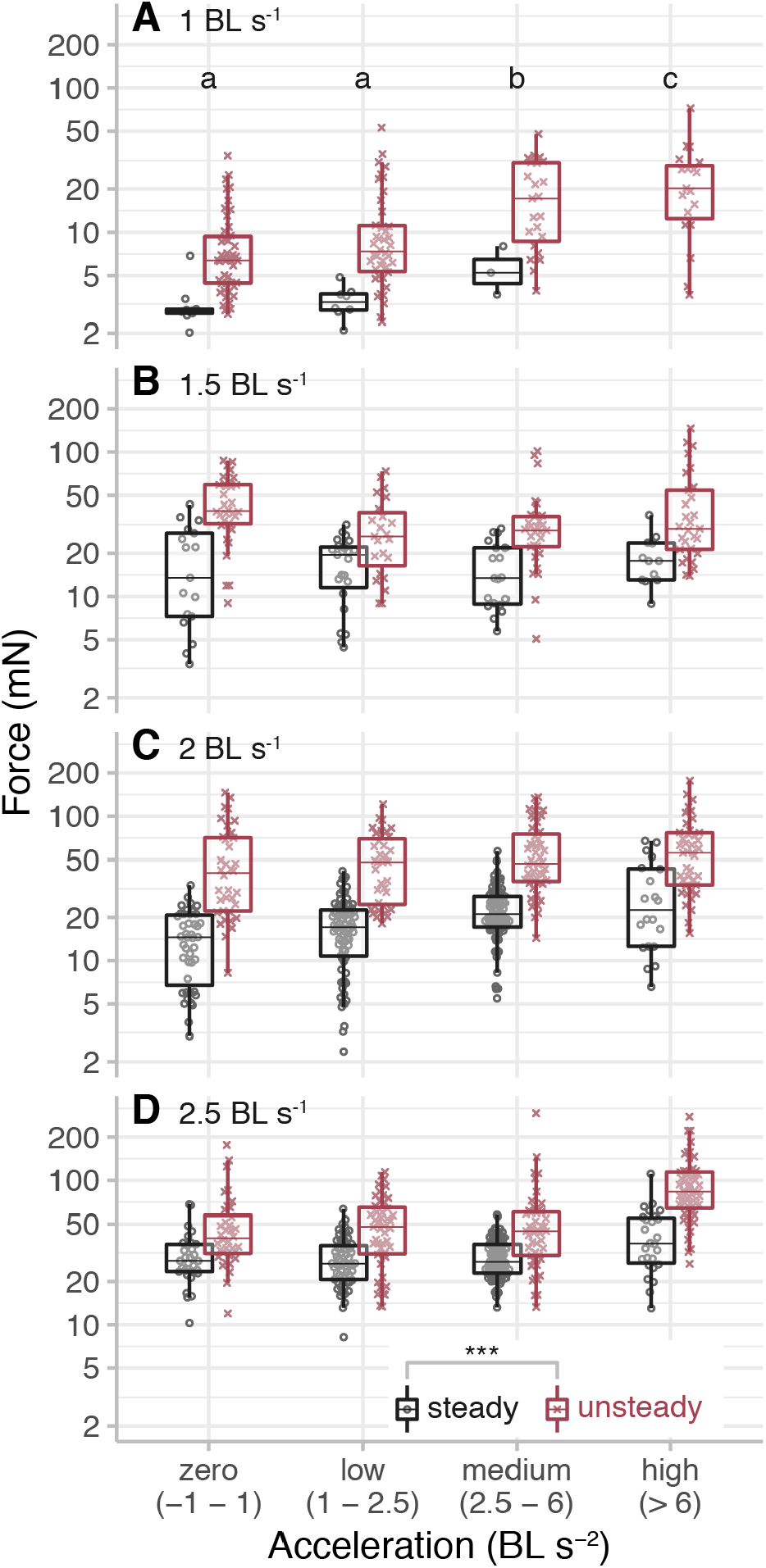
Total force increases in unsteady sequences and with increasing acceleration and swimming speed. Points are jittered to avoid overlap. Letters indicate significant differences among acceleration categories (p < 0.05). ***, p <0.0001.

Based on these results, we can estimate the added mass and drag coefficients using Eq. (1). For the added mass coefficient, we assume that the force to overcome drag is small relative to the acceleration force, except in the zero acceleration bin. Therefore, *C_A_* ≈ *F_total_/ma.* We find that the median added mass coefficient is always higher in unsteady tailbeats, and that it tends to decrease as the acceleration increases (Fig. 6), for example from 1.41 for unsteady, low accelerations to 0.44 for unsteady, high accelerations. It also tends to be higher for accelerations from higher swimming speeds, so that the largest coefficient (2.62) is for low accelerations from 2.5 BL s-1 and the smallest (0.25) is for high accelerations from 1.5 BL s-1.

**Fig. 6:**
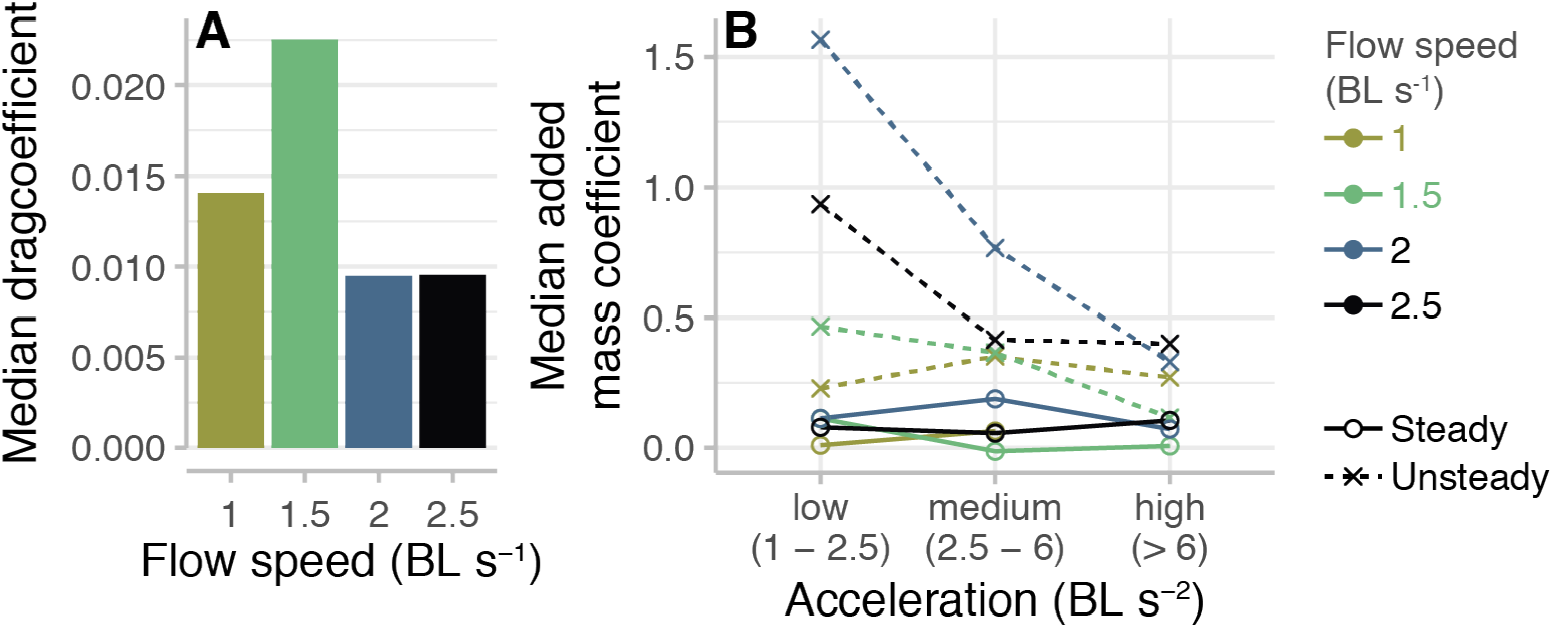
The median added mass coefficient is higher in unsteady sequences and decreases as acceleration increases. The median added mass coefficient *C_A_* is plotted against acceleration groups, where color indicating the flow speed.

Similarly, the drag coefficient can be estimated from the steady tailbeats in the zero acceleration category. In this case, we assume that the force to accelerate is zero, so that 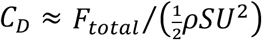. The overall median drag coefficient is 0.011, and it ranges from 0.0095 at 2.5 BL s-1 to 0.023 at 1.5 BL s-1.

The angle of vortex pairs in the wake increases slightly as acceleration increases (Fig. 7). The angle is significantly different among acceleration categories (p = 0.0084; Table 1), but does not change significantly between steady and unsteady sequences (p = 0.0522). Although there is a significant effect of acceleration, the magnitude of the effect is small; the largest difference is between zero and high acceleration, but it is only 4.8±1.3°.

**Fig. 7:**
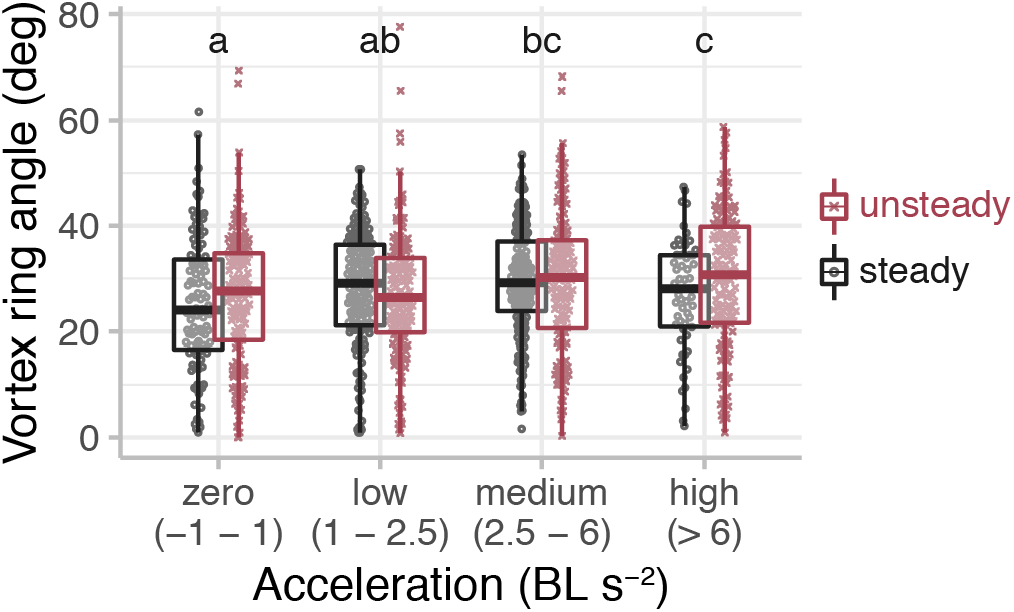
The angle of the vortex ring in the wake increases with increasing acceleration, but is not different among steady and unsteady sequences. The angle of the first vortex pair relative to the swimming direction is plotted against acceleration categories for steady and unsteady sequences. Letters indicate significant differences among acceleration categories.

The vortex ring diameter is significantly smaller for unsteady sequences compared to steady ones (p = 0.0006; Table 1). Fig. 8 shows the horizontal vortex ring diameter *D* normalized relative to its vertical diameter *d*, which we assume is equal to the height of the fish’s tail. This diameter also decreases significantly as flow speed increases (p < 0.0001). A value of *d/D* equal to one indicates a circular ring, and in all cases, the value is significantly different from 1 (p > 0.099), except for the zero acceleration case at 1 BL s-1, in which *d/D* = 1.24±0.06, which is significantly larger than one (p = 0.0293).

**Fig. 8:**
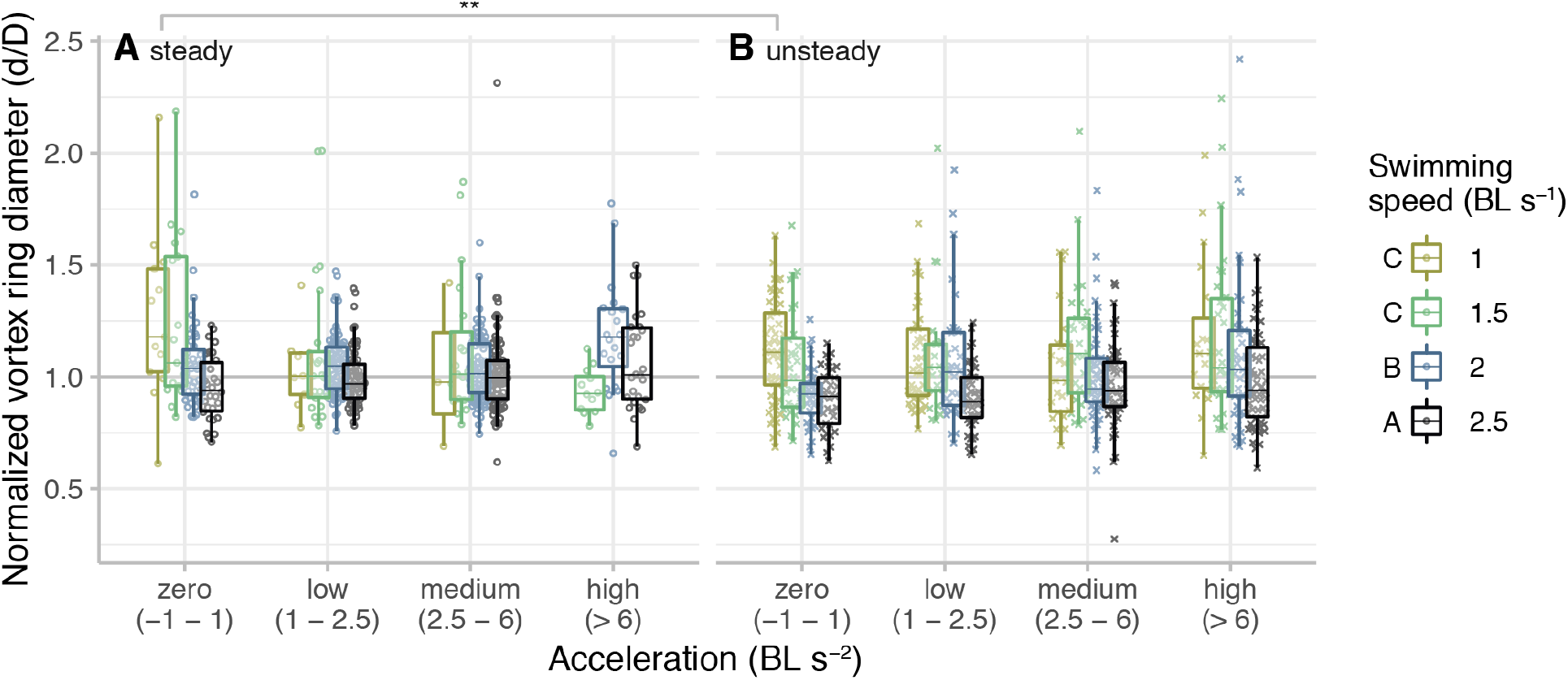
The ratio of the horizontal vortex ring diameter (*D*) to the vertical diameter (*d*) is significantly smaller in unsteady sequences and at higher swimming speeds, but does not change significantly with acceleration. The vortex ring diameter in the horizontal plane *D* is normalized by the tail height *d.* A value of 1 indicates a circular ring. Letters indicate significantly differences in diameter among swimming speeds. **, p < 0.001.

### Kinematic changes during acceleration

Tail beat frequency increases with both flow speed and acceleration, and is higher in unsteady tailbeats compared to steady ones (Fig. 8). Tail beat frequency changes significantly across acceleration categories (p < 0.0001; Table 2) and across flow speeds (p < 0.0001). It is also significantly higher in unsteady tailbeats than in steady ones (p < 0.0001); unsteady tailbeats have a tail beat frequency 1.05±0.51 Hz higher than steady ones, on average. There is also a significant interaction between acceleration and flow speed (p = 0.0002), so that the highest tail beat frequencies occur at 2.5 BL s-1 and high acceleration.

**Table 2.** Mixed model regression results for tail beat frequency and head and tail amplitude.

|  | Frequency<br>(log transformed) |  |  | Head amplitude |  |  | Tail amplitude |  |  |
| --- | --- | --- | --- | --- | --- | --- | --- | --- | --- |
|  | *F* | *df* | *p* | *F* | *df* | *p* | *F* | *df* | *p* |
| Intercept | **402.67** | **1,1096** | **<0.001** | **259.53** | **1,1096** | **<0.001** | **188.09** | **1,1096** | **<0.001** |
| Acceleration categories<sup>1</sup> | **108.74** | **3,1096** | **<0.001** | **17.28** | **3,1096** | **<0.001** | **47.77** | **3,1096** | **<0.001** |
| Tailbeat type<sup>2</sup> | **143.45** | **1,1096** | **<0.001** | **13.49** | **1,1096** | **<0.001** | **114.01** | **1,1096** | **<0.001** |
| Flow speed<sup>3</sup> | **146.21** | **3,1096** | **<0.001** | **17.85** | **3,1096** | **<0.001** | **40.39** | **3,1096** | **<0.001** |
| Tailbeat type ×<br>Acceleration | 0.43 | 3,1096 | 0.734 | 0.58 | 3,1096 | 0.629 | 2.38 | 3,1096 | 0.068 |
| Flow speed ×<br>Acceleration | **3.63** | **9,1096** | **<0.001** | 2.01 | 9,1096 | 0.035 | **4.49** | **9,1096** | **<0.001** |
| Tailbeat type ×<br>Flow speed | **15.98** | **3,1096** | **<0.001** | 2.92 | 3,1096 | 0.033 | **14.38** | **3,1096** | **<0.001** |

<sup>1</sup>zero (-1 to 1 BL s<sup>2</sup>), low (1 – 2.5 BL s<sup>2</sup>), medium (2.5 – 6 BL s<sup>2</sup>), and high (> 6 BL s<sup>2</sup>); <sup>2</sup>steady or unsteady; <sup>3</sup>1.0 BL s<sup>-1</sup>, 1.5 BL s<sup>-1</sup>, 2.0 BL s<sup>-1</sup>, or 2.5 BL s<sup>-1</sup>

Head and tail amplitude are significantly higher during unsteady tailbeats than during steady ones (Fig. 10; p ≤ 0.0003; Table 2). On average, head and tail amplitudes are 0.0037±0.0007 BL and 0.015±0.001 BL higher, respectively, in unsteady tailbeats than steady. Amplitudes are also significantly different among acceleration categories (p < 0.0001 in both cases), but only the high acceleration category has significantly larger amplitudes than the others. They both also increase significantly as flow speed increases (p < 0.0001 in both cases). Tail amplitude is significantly different at each flow speed, while head amplitude at the 1 and 1.5 BL s-1 is significantly different from head amplitude at 2 and 2.5 BL s-1.

The total force is strongly correlated with the frequency and amplitude. The correlation coefficients between log force and frequency, tail amplitude, and head amplitude are 0.74, 0.38, and 0.19. Most of these correlations are related to the changes in vortex circulation, not the changes in vortex diameter. Vortex diameter is best associated with tail amplitude, but the correlation coefficient is only 0.14.

## DISCUSSION

During steady swimming, fish produce a wake that contains regularly spaced vortex pairs (Lauder and Tytell, 2005). The circulation and orientation of these vortices reflect the forces the fish produces for swimming, which include both lateral and axial components. To accelerate, fish must produce more axial force than it would during steady swimming. The axial force could be increased by redirecting the same total force so that it points more posteriorly (Fig. 1B), by increasing the total force output so that the axial component is greater (Fig. 1C), or both. If the angle of the vortex pairs in the wake changes, that would indicate a change in the direction of the force, and if the circulation of the vortices changes, that would indicate a change in the magnitude of the force. Eels change the direction of the force when they accelerate, indicated by a change in the orientation of vortices in their wakes to accelerate (Tytell, 2004a). Since the bluegill has a stiffer body than the eel (Aleyev, 1977), we predicted that it would be less able to curve its body in order to redirect forces and manipulate the locations of vortices in its wake. Instead, we predicted that it would increase the circulation of vortices in the wake, increasing the total force. Our data partially support our hypothesis. The bluegill substantially increased circulation of vortices in the wake (Fig. 5), leading to an increase in total force as acceleration strength increased. It also significantly increased the angle of the vortex pairs, but to a smaller degree (Fig. 7).

Even though bluegill change both the magnitude and direction of force during acceleration, the most important effect is the change in magnitude. The angle of the vortex ring in the wake and the total force it represents allows an estimate the axial force *F_ax_ = F_total_* sin(90 – *θ*).

Axial force increases from 4.2 mN in the steady, zero acceleration case to 21.9 mN in the unsteady, high acceleration case. If the vortex ring angle did not change over this range, the change in total force would still account for 78% of the total increase in axial force. If the vortex ring angle changed, but the total force stayed constant, then the axial force would only increase to 24% of its unsteady, high acceleration value. Thus, as we hypothesized, the primary way the bluegill accelerates is to increase the force it produces, not by redirecting the force or changing the structure of its wake.

This pattern is different from the changes in the wakes of accelerating eels (Tytell, 2004a). Steadily swimming eels produce lateral jets in their wake, with very little downstream momentum, as is required by the zero net force on the body during steady swimming (Tytell and Lauder, 2004). As eels accelerate, they change the structure of their wake, rotating the jets to point backwards, which provides the extra force needed to accelerate (Tytell, 2004a). The eels do not substantially change the circulation of the vortices in the wake.

Carp may accelerate using a combination of these two patterns. Wu et al. (2007) studied burst-and-coast swimming and compared bursts with a single tail flick to one side to those with multiple tail flicks. When bursting with a single tail flick, they change the angle of the vortex pair substantially compared to a burst with multiple tail flicks (Wu et al., 2007). They do not report any data from steady swimming, so it is not possible to ascertain if the vortex circulation was higher during acceleration compared to steady swimming.

The kinematic changes we observed are consistent with those described by Akanyeti et al. (2017), who performed a large survey of acceleration in many species of fishes. They found that tail beat amplitude increased by 34% during acceleration, relative to steady swimming. In our data, the tail beat amplitude during the unsteady, high acceleration case is 33±15% higher than the steady, zero acceleration case (Fig. 10). Akanyeti et al. also reported that tail beat frequency increased with acceleration and with swimming speed, but that acceleration was the stronger effect, the same pattern that we observed (Fig. 9).

**Fig. 9:**
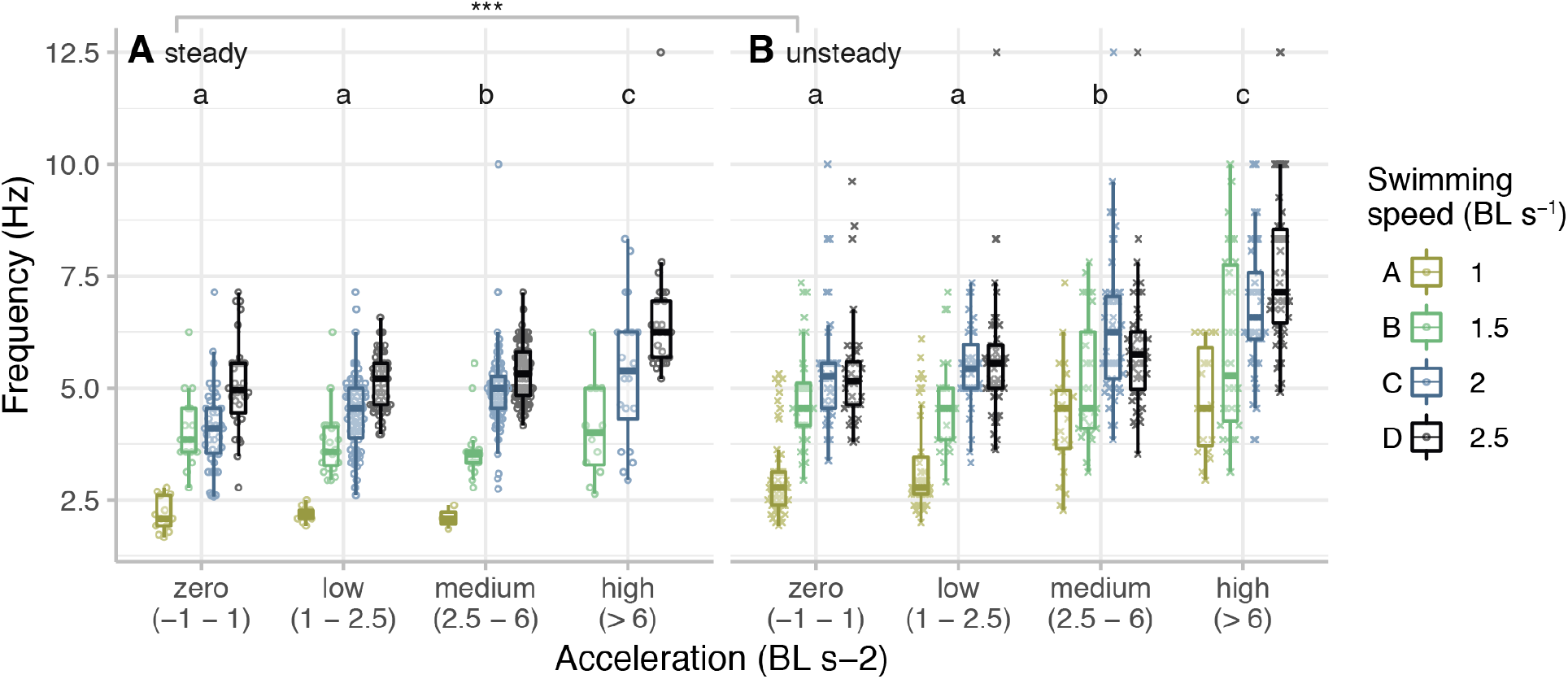
Tail beat frequency is higher in unsteady sequences and increases with both acceleration and swimming speed. Tail beat frequency is plotted against acceleration categories for steady (A) and unsteady (B) sequences. Letters indicate significantly differences in diameter among swimming speeds. ***, p < 0.0001.

**Fig. 10:**
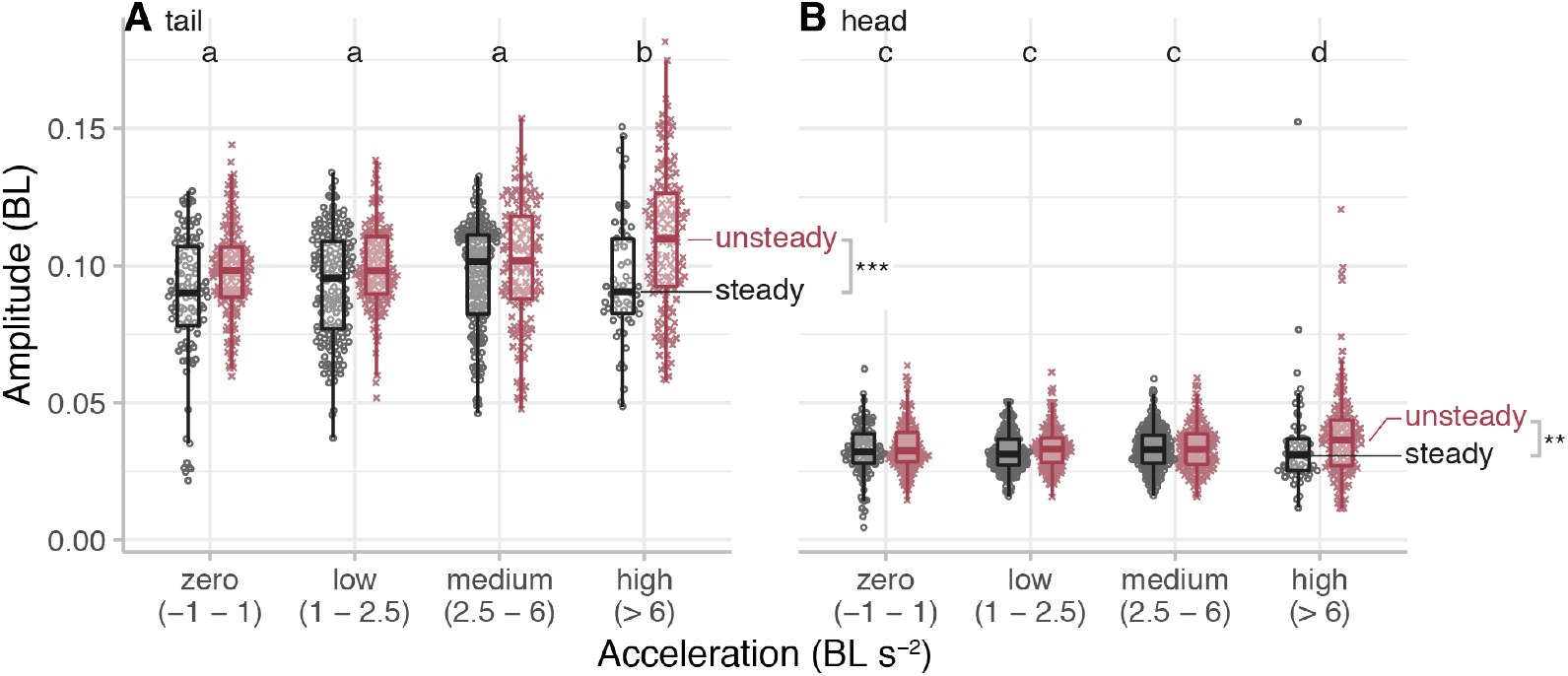
Both head and tail amplitude increase in unsteady sequences and at the highest accelerations. Tail amplitude (A) and head amplitude (B) are plotted against acceleration categories for steady sequences (black) and unsteady sequences (red). Letters indicate significantly differences in amplitudes among acceleration categories. **, p < 0.001; ***, p < 0.0001.

However, our wake flow data are different from those of Akanyeti et al. (2017). They hypothesized that accelerating fishes modulate the vortex ring size and orientation to increase propulsive efficiency, but our data do not support this hypothesis. They performed detailed wake analysis from trout Oncorhyncus mykiss during acceleration and found that vortex ring impulse increased dramatically, the vortex ring jet reoriented substantially more downstream, and the rings became more circular. In our data, we find a similar increase in impulse. For trout, the impulse increased by 3.25 times (based on their Fig. 4C; Akanyeti et al., 2017). We found that the median impulse in the unsteady, high acceleration case was 2.1 times the value in the steady, zero acceleration case. We did not find a substantial change in vortex angle (only 4.8±1.3°), while Akanyeti reported a change of 28° (based on their Fig. 4C; Akanyeti et al., 2017). We found no support for the idea that vortex rings become more circular during acceleration. Bluegill vortex rings are not significantly different from circular at nearly all combinations of flow speed and acceleration. Trout produce oval-shaped rings during steady swimming (*d/D <*
1) and decrease the horizontal diameter *D* during acceleration, so that *d/D* becomes closer to 1. Bluegill show the opposite pattern: *d/D* starts out higher during steady tailbeats and decreases during unsteady ones.

### Steady and unsteady tailbeats

Even in trials in which the fish swam steadily, we found a range of acceleration magnitudes (Fig. 3). As part of the experimental procedure, we performed trials in which we elicited accelerations, and performed other trials in which we worked with the fish until it swam steadily for at least 5 full tail beats. Steady swimming was straightforward to assess because the fish were swimming against a steady flow in a flow tank. When the fish matched the flow speed, the image in the video would remain in the same place over several tail beats, moving less than 2% of its body length within the flow tank. For steady and unsteady tailbeats, we measured the acceleration using the IMU. Every tail beat produces a small acceleration and a small deceleration, even when the fish is swimming steadily on average (Borazjani, 2015; Plew et al., 2007; Tytell, 2007; Wen and Lauder, 2013; Xiong and Lauder, 2014), and the range of accelerations increases at higher steady swimming speeds (Plew et al., 2007; Xiong and Lauder, 2014). Most of the steady tailbeats had zero or low acceleration (Fig. 3), but there was still a range of acceleration magnitudes, and the range of accelerations increased as the swimming speed increased, similar to what Xiong and Lauder (2014) reported. In unsteady tailbeats, however, the range of accelerations was very comparable across swimming speeds.

The primary different between steady and unsteady tailbeats is that, in unsteady tailbeats, the fish maintains an acceleration over several tail beats. A steady trial may have a tail beat with a strong forward acceleration, but it is followed by a tail beat with a similar deceleration, so that the overall speed does not change. In unsteady tailbeats, a strong acceleration in one tail beat is sustained over several more, so that the overall speed increases.

Because of this range of accelerations, we could compare steady and unsteady tailbeats with the same acceleration magnitude. Surprisingly, the fish produced substantially higher forces during unsteady tailbeats than during steady ones, even at the same acceleration (Fig. 5). How could this be possible? We suggest that the extra force is needed to overcome fluid dynamic added mass (Faber, 1995), and that the added mass coefficient increases when both tail beat amplitude and frequency increase. When a fish is actively trying to accelerate, it must overcome the fluid dynamic acceleration reaction, also called added mass (Daniel, 1984; Faber, 1995). To accelerate in a fluid, the force required is

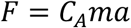

where *m* is the mass, *ρ* is the fluid density, *V* is the volume of the body, *a* is the acceleration, and *C_A_* is the added mass coefficient, which is typically 1.0 or less for streamlined bodies (Daniel, 1984). An intuitive explanation for added mass is that an accelerating body in a fluid must also accelerate some of the fluid around it. Based on our data, we estimated added mass coefficients for the bluegill (Fig. 6B). These coefficients were always higher for unsteady tailbeats, in which acceleration was sustained over several tailbeats, compared to steady tailbeats, and they decreased with increasing acceleration.

We suggest that the reason for the increase in force in unsteady tailbeats is that higher tailbeat amplitudes and frequencies increase the added mass coefficient. If the body amplitude is higher, then the amount of fluid that must be accelerated with the body is also higher. At higher tail beat frequencies, the side-to-side acceleration of the body is also higher. Each time the tail beats, the fish accelerates forward (Tytell, 2007; Xiong and Lauder, 2014). At a higher tail beat frequency, these small accelerations occur more frequently, potentially leading to a greater impact from the acceleration reaction over the course of multiple tailbeats.

We found that head and tail amplitude both increased by about 12-32% in unsteady tailbeats, relative to steady (Fig. 9). Frequency increased by 1.05±0.51 Hz in unsteady tailbeats. Both of these increases could increase the added mass. Tytell (2004a) estimated the added mass coefficient for accelerating eels and found that it was as high as 2.8 during accelerations. Similarly, Wu et al. (2007) estimated the drag coefficient on carp during acceleration and found that it was about four times higher than the drag during gliding. However, they assumed a constant added mass coefficient. Their results could also be explained by an increase in added mass during acceleration, as they acknowledge (Wu et al., 2007).

### Kinematics during accelerations

Relatively few studies have quantified how kinematics change as a function of the magnitude of an acceleration. Like the current results, Tytell (2004a) found that eels accelerate by increasing both head and tail amplitude proportionally to acceleration. Bluegill also increase amplitude during accelerations, compared to steady swimming, but the increase is not proportional to the acceleration (Fig. 9). Eels also increase tail beat frequency during acceleration (Tytell, 2004a), much like bluegill in our study.

Acceleration performance is related to increases in amplitude across the whole body (Akanyeti et al., 2017). Across a wide range of species, Akanyeti et al. (2017) found that undulatory amplitudes increased by about 34% during acceleration, which is very close to the 33% that we found in unsteady, high accelerations. Similarly, Wen et al. (2012) and Borazjani and Sotiropoulos (2010) studied the swimming performance of robotic and computational models as they accelerated from rest to a steady state, comparing the performance of anguilliform (eel-like) and carangiform (mackerel-like) kinematics. For the same tail amplitude, anguilliform swimmers have higher anterior body amplitude, and both studies found that these kinematics cause more rapid initial acceleration, even though their final swimming speed is lower (Borazjani and Sotiropoulos, 2010; Wen et al., 2012). These models thus suggest that carangiform swimmers like bluegill can accelerate faster by adopting a more eel-like swimming mode with greater head amplitude, which is what our data show (Fig. 10).

We also found that frequency increases proportionally to both swimming speed and acceleration (Fig. 9), but that the increase for acceleration is larger than the increase for swimming speed. This is similar to the pattern that Akanyeti et al. (2017) reported for trout.

### Contributions of other fins to acceleration

In this study, we only examined the contribution of the caudal fin to thrust, but, clearly, other fins also contribute. Although we did not quantify it, one can observe from our videos that bluegill tend to beat their pectoral fins at the beginning of an acceleration. Thus, the force we measured off of the caudal fin is not the total force on the entire body. The role of the pectoral fin should diminish as flow speed increases. However, as the steady swimming speed increases, pectoral fin forces are directed more laterally, so their contribution to thrust might decrease (Drucker and Lauder, 2000). The angle between the force vector and the swimming direction increases to around 86° as flow speeds increase past 1.5 BL s^−1^ (Fish and Lauder, 2006). It is not known whether the pectoral fins re-orient their forces to produce more thrust during acceleration. If the pattern seen during steady locomotion remains during accelerations, then the pectoral fins should produce very little axial force in comparison to the caudal fin.

The dorsal and anal fin may also produce significant thrust forces during accelerations. These fins have been shown to produce significant amounts of force once flow speed surpasses 1.1 BL s^−1^. The dorsal fin alone can produce 12% of total thrust during steady swimming (Drucker and Lauder, 2001). The anal fin may have similar thrust patterns (Tytell, 2006). It seems likely that these median fins are also important for thrust during acceleration.

Individual fish may partition forces among their fins differently. We observed that certain fish used their pectoral and caudal fins at different times. Some of the fish used their pectoral fins very infrequently, and others did not. If these other fins produce different amounts of force in different individuals, it could explain the large variation of force produced by the caudal fin for a given acceleration (Fig. 5). We found that individuals varied significantly in how much force they produced and how the force changed from steady to unsteady tailbeats. This variation could be a result of individuals relying on different fins for the same thrust requirements.

### Efficiency and stability during acceleration

We found that bluegill accelerate primarily by increasing the total force in their wake and
only secondarily by changing the angle of vortex pairs (Fig. 7). However, altering the angle would be a more energetically efficient way to accelerate. If bluegill could simply reorient the vortex rings in the wake so that more of the force was directed backwards, then they would not have to expend more energy to create stronger vortices with higher circulation. To change the vortex ring orientation, the bluegill would have to change the lateral spacing of vortices relative to the distance it travels forward in a single tail beat (called the “stride length”) (Videler, 1993). In Fig. 1B, the lateral spacing is the same as Fig. 1A, but the axial distance between vortices is less. Alternatively, bluegill could increase the lateral spacing of vortices while keeping the stride length constant.

However, physics may constrain how much the bluegill can change the geometry of its wake. It may not be able to change its stride length independently of the spacing of vortices in its wake. If it were able to increase the angle of vortices in its wake by decreasing the stride length, then more of the vortex momentum would be aligned in the axial direction, which would tend to increase the stride length. Similarly, if it increased the lateral spacing of vortices while keeping stride length constant, then the central jet between them would be larger and would contain more momentum, which would tend to increase the stride length. Additionally, the bluegill’s body may not be flexible enough to manipulate its wake structure, like the eel does (Tytell, 2004a). In
particular, it would have to flex its tail more to increase the lateral spacing of vortices. Thus, the bluegill’s body mechanics and the physics of propulsion may limit how much it can alter its wake for acceleration.

Even if the bluegill was capable of reorienting vortices in its wake, doing so might sacrifice stability. Reorientation would result in more axial force from the caudal fin, but would also result in smaller lateral forces. At higher flow speeds, bluegills increase lateral forces from the pectoral fins to increase stability during steady swimming (Fish and Lauder, 2006). Similarly, lateral forces from the caudal fin could also help to stabilize the fish. Lateral stabilizing forces may be particularly important during acceleration, because the movement is inherently unstable. The caudal fin produces a large forward force, but it is located behind the center of mass. Much like backing up a car with a trailer attached, this situation represents an unstable equilibrium. Active lateral stabilization may therefore be particularly important during acceleration.

## Conclusions

Bluegill sunfish accelerate by increasing the total amount of force produced during each tail beat, but do not substantially redirect the force produced. This process increases the total amount of axial force, allowing for acceleration, but does not lead to a dramatic reconfiguration of the wake structure, like for acceleration by eels (Tytell, 2004a). The bluegill may be constrained by its relatively stiff body, as well as the physics of propulsion in a fluid, to produce the same sort of wake during steady swimming and acceleration. Similarly, the consistent lateral and axial forces shed by the tail may be necessary in order to stabilize the moving fish. Surprisingly, for the same magnitude acceleration, we found that bluegill produce much lower forces during a single steady tailbeat than within an acceleration sequence that lasts for several tailbeats. We attribute this difference to the increase in amplitude during sustained accelerations, which is required to produce higher forces, but also increases the added mass coefficient on the fish.

## ACKNOWLEDGEMENTS

The authors thank Alexandra Boden for help in running the experiments and Vishesh Vikas with help setting up the IMUs.

## DATA AVAILABILITY

Raw video data are available from ZMAPortal (https://zmaportal.org) study ZMA18. Notes, processed data, and analysis notebooks are available at doi:10.6070/H4KS6Q5V (http://dx.doi.org/10.6070/H4KS6Q5V).

## AUTHOR CONTRIBUTIONS

T.N.W, M.A.B.S, and E.D.T. designed the study. T.N.W. performed the experiments. E.D.T. wrote software to process the data. T.N.W. and E.D.T. analyzed the data. T.N.W. wrote the initial draft, and M.A.B.S. and E.D.T. edited the draft. E.D.T. acquired funding for the project.

---

**Movie S1. Example video, flow patterns, and kinematic variables during steady swimming at 1.5 BL s^−1^**. The top panel shows video, filmed from below, with flow vectors in color. The color indicates vorticity. The bottom two panels show the fish’s relative speed within the flow tank (white solid line), the acceleration (brown dotted line), the position of the tail (blue dashed line), and the absolute value of the circulation of vortices shed into the wake (red circles). The current time within the video is shown with a dashed white line, and the time that corresponds to Fig. 4A is shown with a dotted white line. The video is slowed by 10 times.

**Movie S2. Example video, flow patterns, and kinematic variables during a medium acceleration starting at 1.5 BL s^−1^.** The top panel shows video, filmed from below, with flow vectors in color. The color indicates vorticity. The bottom two panels show the fish’s relative speed within the flow tank (white solid line), the acceleration (brown dotted line), the position of the tail (blue dashed line), and the absolute value of the circulation of vortices shed into the wake (red circles). The current time within the video is shown with a dashed white line, and the time that corresponds to Fig. 4B is shown with a dotted white line. The video is slowed by 10 times.

**Movie S3. Example video, flow patterns, and kinematic variables during a high acceleration starting at 1.5 BL s^−1^.** The top panel shows video, filmed from below, with flow vectors in color. The color indicates vorticity. The bottom two panels show the fish’s relative speed within the flow tank (white solid line), the acceleration (brown dotted line), the position of the tail (blue dashed line), and the absolute value of the circulation of vortices shed into the wake (red circles). The current time within the video is shown with a dashed white line, and the time that corresponds to Fig. 4C is shown with a dotted white line. The video is slowed by 10 times.

